# Divergent regulatory impacts of endogenous siRNAs on host mRNAs in testis of closely related species

**DOI:** 10.64898/2026.05.18.725984

**Authors:** Mina Zamani, Taiya Jarva, Henry Moukheiber, Ava Kloss-Schmidt, Nandagopal Ajaykumar, Ching-Jung Lin, Berta Chalouh-Hara, Avishye Moskowitz, Josefa Steinhauer, Eric C. Lai, Ildar Gainetdinov

**Author notes:** These authors contributed equally (Mina Zamani, Taiya Jarva).

## Abstract

Invertebrates use RNAi to fight viruses that replicate via dsRNA intermediates, which serve as precursors for producing anti-viral siRNAs. In addition, siRNAs are produced from a broad range of endogenous dsRNAs in *Drosophila melanogaster*. However, beyond the impacts of siRNAs derived from a handful of hairpin RNAs, the regulatory potential of most endo-siRNAs has been unknown. Here, we report that RNAi is far more potent in the close sister species *Drosophila simulans*. Consequently, endo-siRNAs repress less than a dozen transcripts in *D. melanogaster*, but hundreds of mRNAs in *D. simulans* testis. These regulatory interactions occur in cis (between pairs of bidirectionally transcribed loci), as well as in trans (between genomically unlinked siRNA-target pairs). We establish the molecular determinants of productive gene repression by siRNAs *in vivo*, including cleavage in trans via ≥16-nt contiguous complementarity. Our data indicate that *D. simulans* spermatogenesis requires repression of numerous host mRNAs by endo-siRNAs. More generally, we reveal unexpectedly fast-evolving and broad regulatory impacts of endogenous RNAi in the male germline.

## INTRODUCTION

In eukaryotes, small noncoding RNAs direct Argonaute proteins to regulate complementary targets.^1^ Animals produce three major classes of small RNAs that mediate repression via Argonaute effectors. MicroRNAs (miRNAs) repress host mRNAs,^2,3^ PIWI-interacting RNAs (piRNAs) silence transposons,^4,5^ and small interfering RNAs (siRNAs) in the RNA interference (RNAi) pathway are essential for antiviral defense.^6–9^ RNAi uses double-stranded RNAs (dsRNAs) as substrates and thus efficiently represses viruses that replicate via dsRNA intermediates.^6–8^ While RNAi has been directly shown to be essential for antiviral defense in several invertebrate species, vertebrates generally rely on the innate immune pathway to fight RNA viruses.^10^

The RNAi pathway also generates endogenous siRNAs (endo-siRNAs) from host transcripts that form dsRNA, in both flies^11–17^ and mice.^18–20^ Vertebrates generally lack endo-siRNAs. The oocytes of mice and rats (Murinae) are an exception.^18,19^ Murinae regained the ability to produce endo-siRNAs because a transposon insertion in Dicer generates an oocyte-specific isoform encoding Dicer^O^, which lacks the amino-terminal domain that normally blocks the use of long dsRNA substrates.^21^

While the number of endo-siRNA-producing loci in *Drosophila melanogaster* is greater than the number of miRNA loci,^22^ there is limited evidence that they influence gene expression. *Drosophila* endo-siRNAs with known functions derive from hpRNAs and repress selfish sex ratio distorters.^20,23–25^ A subset of transposon transcripts is modestly upregulated in fly RNAi mutants.^11–13,16,26^ However, evidence for the repression of host genes by *Drosophila* endo-siRNAs is sparse or equivocal, despite the fact that all cis-NAT siRNAs in principle have a perfectly complementary target.^15,18,19,27^

Here, we report that endo-siRNAs have distinct regulatory impacts in the testis of the closely related species *Drosophila melanogaster and Drosophila simulans*, separated by just five million years of evolution. siRNAs in *D. melanogaster* are present at modest levels and repress only a handful of host mRNAs. In *D. simulans*, the greater efficiency of siRNA biogenesis results in the higher abundance of siRNAs and permits repression of numerous endogenous transcripts. First, *D. simulans* siRNAs impact expression of hundreds of host cis-NAT mRNAs *in cis*. Second, our analyses define the molecular requirements for siRNA cleavage in vivo and show that abundant hpRNA-derived siRNAs repress dozens of imperfectly paired targets *in trans*. We propose that regulation of many host mRNAs by endo-siRNAs is required for successful male germline development in *D. simulans*. Together, these data reveal an unexpectedly broad regulatory influence of endogenous RNAi and establish in vivo parameters of biogenesis and gene repression by endo-siRNAs.

## RESULTS

### siRNA biogenesis is highly potent in *Drosophila simulans* testis

Production of siRNAs begins with Dicer-2-catalyzed cleavage of long dsRNAs at regular, 21-nt intervals.^28–31^ Therefore, the vast majority of siRNAs generated from perfectly double-stranded RNA are 21 nt long. However, some 22–23-nt siRNA species are produced from hpRNAs that contain 1–2 bulged unpaired nucleotides. Moreover, a fraction of siRNA 3′ ends are shortened (trimmed) by exonucleases or extended (tailed) by terminal nucleotidyl transferases.^32,33^ We thus considered 20–24-nt small RNA reads with the same 5′ prefix as a single small RNA species. To identify *bona fide* endogenous siRNAs, we searched for 20–24-nt small RNAs present in the control (≥1 ppm) and reduced ≥10-fold or undetectable in *dcr-2* mutant *Drosophila* testis (Figure 1A). No small RNAs derived from arthropod viruses were detected in the control or *dcr-2* testis.

**Figure 1.**
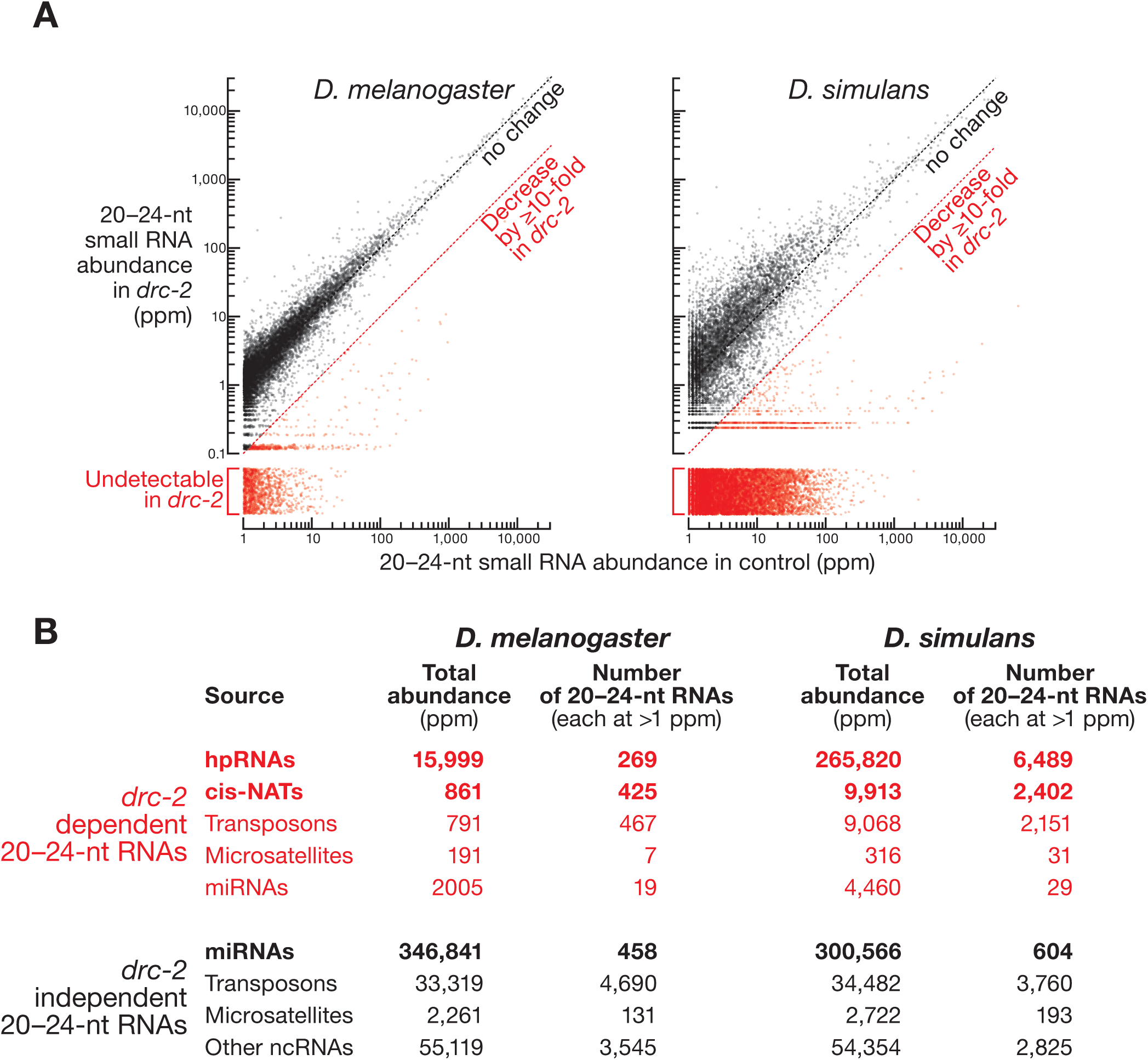
siRNAs are tenfold more abundant in *D. simulans* than *D. melanogaster* testis. (A) Mean abundance of ≥1 ppm, 20–24-nt small RNAs in control and *dcr2* testis in *D. melanogaster* (*n* = 3) and *D. simulans* (*n* = 2). Abundance of 20–24-nt small RNAs with the same 5′ prefix was summed. *dcr2*-dependent siRNAs are defined as decreased by ≥10-fold or undetectable in *dcr2* testis and are shown in red; all other small RNAs are assigned as *dcr2*-independent and are shown in black. (B) Number and total abundance of ≥1 ppm small RNAs mapping to distinct types of genomic loci. *dcr2*-dependent and *dcr2*-independent small RNAs are defined in (A).

Strikingly, the total abundance and number of distinct siRNA sequences were both tenfold greater in *D. simulans* compared to *D. melanogaster* testis: ∼290,000 ppm and ∼11,300 species (each at ≥1 ppm) in *D. simulans* vs ∼20,000 ppm and ∼1,200 species (each at ≥1 ppm) in *D. melanogaster* (Figures 1A and 1B). The increased total siRNA abundance in *D. simulans* could not be attributed to a single type of genomic source. On the contrary, all major classes of siRNA-producing loci (cis-NATs, hpRNAs, transposons) generated a larger number of more abundant siRNA species in *D. simulans* testis (Figure 1B). Repeating analyses for 21-nt small RNAs alone produces virtually indistinguishable results. Our genetic data concur with the recently reported high abundance of siRNAs in wild-type *D. simulans* testis.^34^

*dcr-2*, *loqs*, and *r2d2* mRNAs were expressed at ∼1.5–2.3-fold greater levels in *D. simulans* than in *D. melanogaster* testis, possibly contributing to the more efficient siRNA biogenesis in *D. simulans* (Table S1). Potential differences in the enzymatic properties of the two Dicer-2 homologs could also explain the higher efficiency of RNAi in *D. simulans*. For example, 238 of 1,722 (∼14%) amino acids are different between *D. simulans* and *D. melanogaster* Dicer-2 enzymes. Of the 238 divergent residues, 12 were previously shown to be important for the helicase and nuclease activities of *D. melanogaster* Dicer-2 as well as its interactions with the co-factor protein Loqs (refs.^35,36^; Table S2).

### Molecular determinants of siRNA loading and maturation in *D. simulans*

The high steady-state levels of siRNAs in *D. simulans* afforded an opportunity to probe siRNA biogenesis determinants that were previously established using in vitro experiments and analyses of the limited number of endo-siRNAs in *D. melanogaster*.^37–42^ Fly Dicer-1 and Dicer-2 enzymes produce miRNA and siRNA duplexes, respectively. The double-stranded precursors are then sorted between Ago1 and Ago2 based on the duplex structure.^38^ miRNA precursors contain central mismatches that favor binding to Ago1 and prevent loading in Ago2. By contrast, perfectly complementary siRNA duplexes are overwhelmingly loaded in Ago2 and rarely bind Ago1.^38–41^ Release of the passenger strand from the duplex concludes the formation of Argonaute silencing complex. The strand that remains associated with the Ago—the guide strand—directs Argonaute to complementary targets.

Release and decay of the passenger strand result in its lower abundance, compared to that of the guide strand. Such a bias in guide vs passenger abundance is the hallmark of miRNA and siRNA biogenesis, except for the cases when both strands in a duplex are loaded equally efficiently. Consistent with the idea that perfectly paired siRNA duplexes can only load and mature in Ago2,^38^ our data show that siRNA precursors persist as double-stranded intermediates in *ago2* mutants. When normalized to miRNAs, total siRNA abundance increased ∼1.9-fold in *ago2* testis, compared to control (Figure S1A). However, the strong guide vs passenger abundance bias present in the control was lost in *ago2* testis (Figures 2A and S1B). Double-stranded siRNA intermediates in *ago2* mutants either remain unloaded or, even if loaded in Ago1,^43^ fail to remove the passenger stand.^38,42^ Supporting the idea that siRNA duplexes fail to mature in the absence of Ago2, siRNAs in *ago2* mutant testis were incapable of silencing seed-matched (Figures S1C and S1D) or perfectly complementary targets (see below).

**Figure 2.**
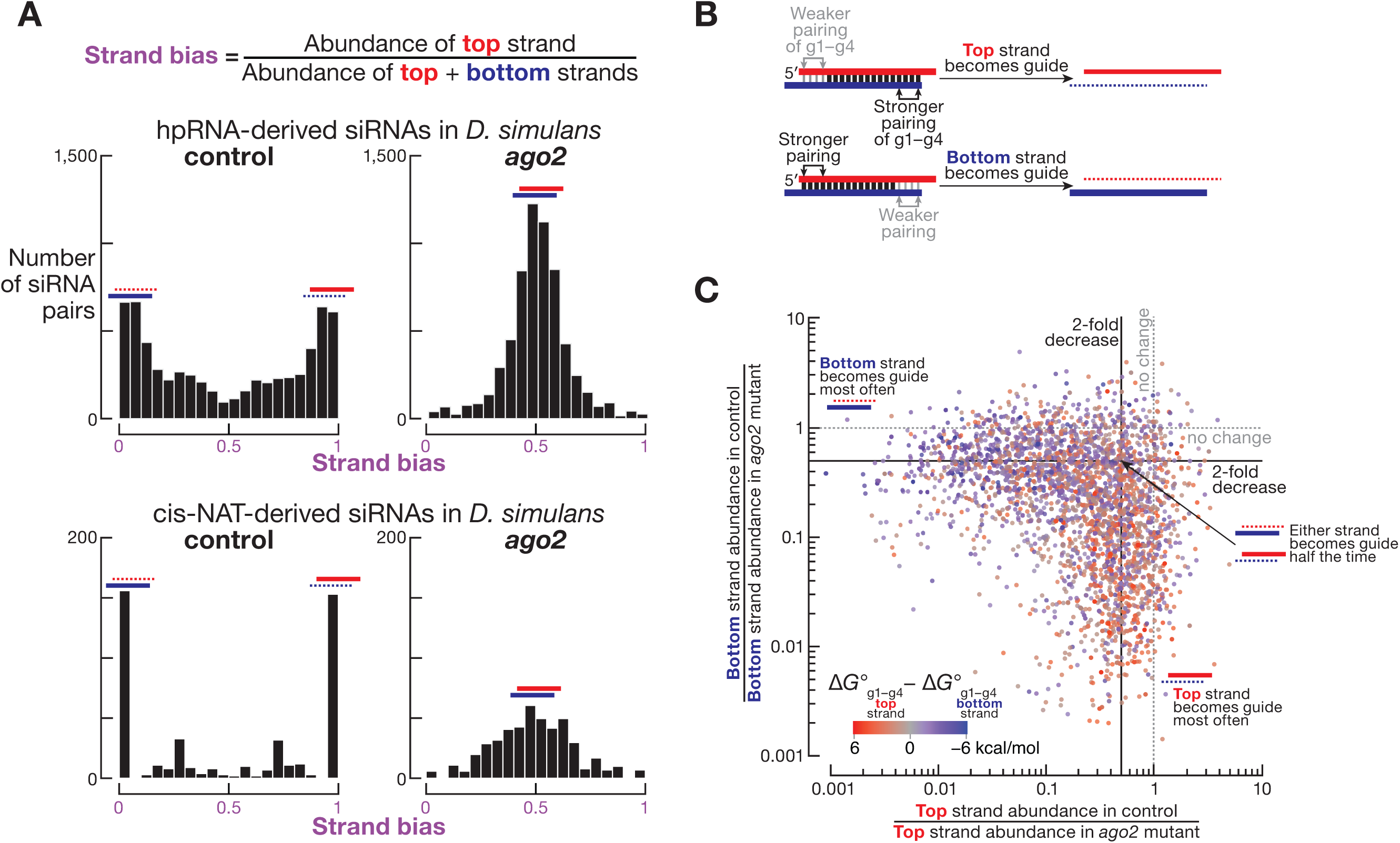
Duplex structure and thermodynamic properties drive siRNA biogenesis in vivo. (A) Mean (*n* = 2) strand bias was calculated as the abundance of top strand divided by the sum of abundances of top and bottom strands. Data are from control and *ago2* testis for perfectly complementary duplexes derived from hpRNA and cis-NAT loci in *D. simulans*. (B) In each siRNA duplex, strand with weak pairing of 5′ terminal nucleotides often becomes the guide. (C) Change in abundance between control and *ago2* mutants for top and bottom strand in each siRNA duplex. Data are colored by the difference in predicted pairing energy of nucleotides g1–g4 of top and bottom strands.

Ago2-bound siRNAs were reported to frequently begin with cytidine (C) or uridine (U) in *D. melanogaster*.^41,42^ However, our analyses of *D. simulans* siRNAs showed only a ∼1.25-fold enrichment for 5′ terminal U and C (U: ∼37.2% in guide strands in wild-type control vs ∼29.8% in both strands of siRNAs duplexes in *ago2* mutants; C: ∼24.4% vs ∼19.5%; Figure S1E). Such modest an influence of 5′ nucleotide identity on loading efficiency in *D. simulans* Ago2 suggests that siRNA biogenesis is not restricted to a subset of substrates.

The thermodynamic properties of an siRNA determine which of the two strands becomes guide or passenger. The guide typically derives from the strand whose 5′ bases are less stably paired in the siRNA duplex (refs.^44,45^; Figure 2B). Consistent with such asymmetry, the difference in the pairing energy of nucleotides g1–g4 between top and bottom strands predicted well which of the two strands became guide or passenger (Figure 2C).

Our analyses also showed that virtually all siRNA duplexes are used to produce siRNA-guided Ago2. When comparing the abundance of strands in siRNA duplexes before maturation (in *ago2* mutants) with that of the mature Ago2-loaded siRNAs (in control testis), we did not find duplexes for which both strands greatly decreased in control tissues (Figure 2C). These data suggest that at least one strand is efficiently loaded when Ago2 is present. We conclude that the duplex structure and its thermodynamic properties, but not the identity of 5′ terminal nucleotide, drive siRNA loading or maturation in *D. simulans*.

### cis-NAT siRNA abundance is dictated by substrate dsRNA levels

Antisense transcripts arising from the same genomic locus (cis-NATs) form dsRNAs that are readily cleaved by Dicer-2 to initiate siRNA biogenesis. The most frequent sources of cis-NAT siRNAs are overlapping 3′ UTRs of convergently transcribed genes. We identified 1,003 syntenic pairs of genes whose 3′ UTRs overlapped both in *D. melanogaster* and in *D. simulans*. Of these, 236 gene pairs produced cis-NAT siRNAs in both species, while 111 generated siRNAs only in *D. melanogaster* and 260 only in *D. simulans* (Table S3).

Both in *D. melanogaster* and *D. simulans*, the abundance of cis-NAT siRNAs was determined by the amount of dsRNA substrate. In theory, the formation of dsRNAs for each gene pair is limited by the abundance of the transcript that is expressed at the lower level. Indeed, we observed a stronger correlation between siRNA levels and the abundance of the mRNA expressed at the lower level in each cis-NAT pair (*D. melanogaster*: Spearman’s ρ = 0.6, *p* = 0; *D. simulans*: ρ = 0.6, *p* = 0; Figure S2A). By contrast, siRNA abundance was less correlated with the steady-state level of the more abundant transcript in a gene pair (*D. melanogaster*: ρ = 0.46, *p* = 0; *D. simulans*: ρ = 0.29, *p* = 0; Figure S2B).

cis-NAT transcript pairs that did not produce siRNAs were typically poorly expressed or undetectable. For example, the median abundance of mRNAs annotated as convergently transcribed but that did not make siRNAs was lower compared to the median abundance of transcripts that generated siRNAs (7-fold in *D. melanogaster*, *p* = 9 × 10^−34^; 24-fold in *D. simulans*, *p* = 2 × 10^−62^; Figure S2C).

Consistent with the more efficient siRNA biogenesis in *D. simulans*, for the 236 gene pairs that made siRNAs in both species, more siRNAs were produced in *D. simulans* compared to its homologous gene pair in *D. melanogaster*, even when normalized to mRNA abundance (median: 3.5-fold more siRNAs in *D. simulans* than in *D. melanogaster*; interquartile range [IQR]: 1.6–6.5-fold; Figure 3A). Together, these data show that dsRNA abundance and the species-specific efficiency of RNAi are the main determinants of cis-NAT siRNA biogenesis.

**Figure 3.**
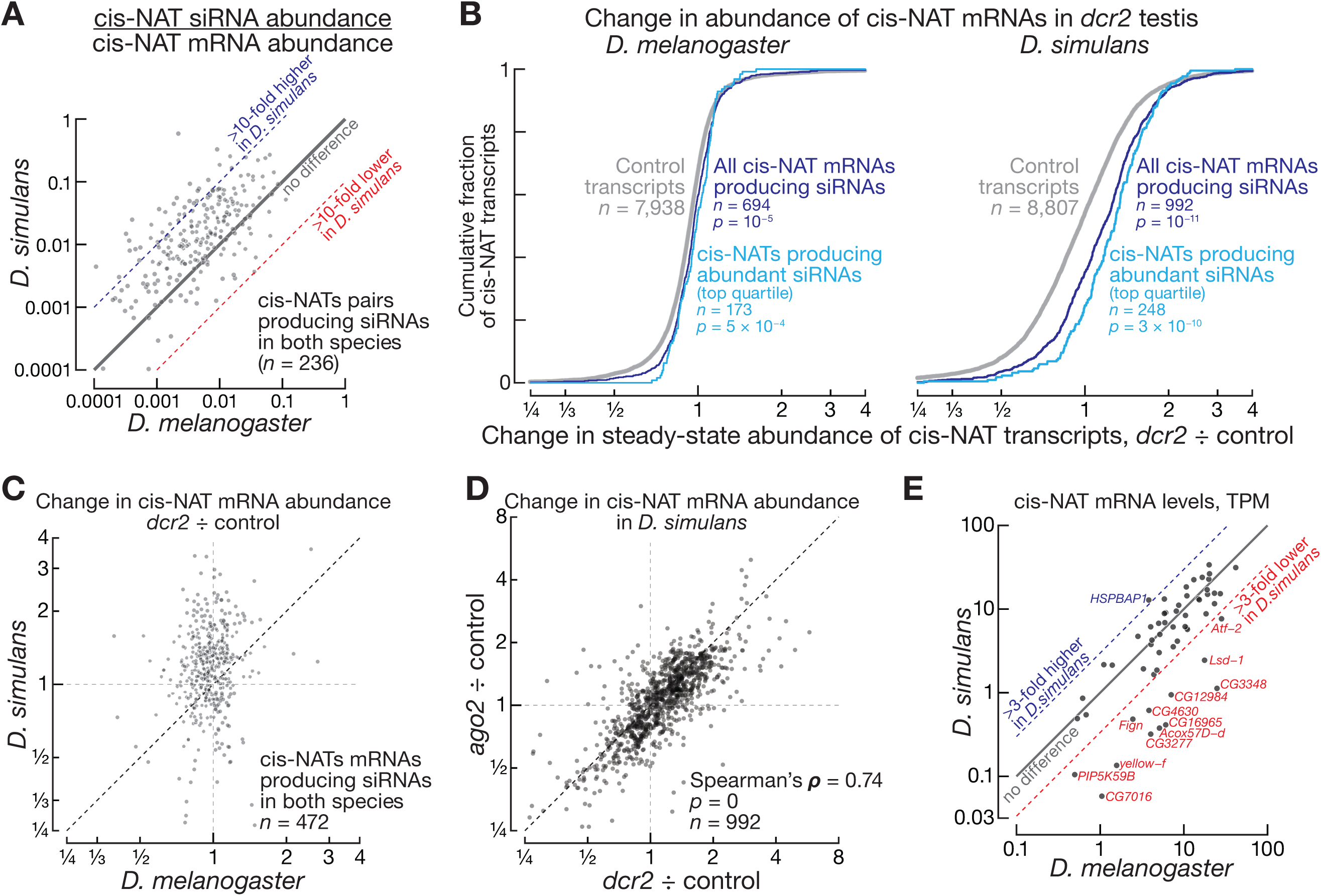
RNAi represses hundreds of cis-NAT mRNAs in *D. simulans* testis. (A) Mean abundance of cis-NAT siRNAs normalized to transcript abundance in *D. melanogaster* (*n* = 3) and *D. simulans* (*n* = 2). Data are for cis-NAT genes that produce siRNAs in both species (Table S3). (B) Change in steady-state abundance of cis-NAT mRNAs in *dcr2* vs control testis in *D. melanogaster* (*n* = 3) and *D. simulans* (*n* = 2). Data are for cis-NAT siRNA-producing genes only (Table S3). *P* values for two-tailed Kolmogorov-Smirnov test are shown. (C) Comparison of change in abundance in *dcr2* vs control testis for cis-NAT mRNAs in *D. melanogaster* (*n* = 3) and *D. simulans* (*n* = 2). Data are for cis-NAT genes that produce siRNAs in both species (Table S3). (D) Comparison of change in abundance for cis-NAT mRNAs between *dcr2* (*n* = 2) and *ago2* (*n* = 2) vs control testis in *D. simulans*. Data are for all cis-NAT genes that produce siRNAs in *D. simulans* (Table S3). Rank-order Spearman’s correlation coefficient and *p* value are shown. (E) Mean steady-state abundance of cis-NAT mRNAs in *D. melanogaster* (*n* = 3) and *D. simulans* (*n* = 2). Data are for 58 cis-NAT genes that are efficiently repressed by RNAi in *D. simulans* (i.e., derepressed by ≥2-fold in *dcr2* mutants; Table S3).

### Ago2 represses hundreds of 3′ UTR cis-NAT mRNAs in *D. simulans* testis

Genes arranged in cis-NAT pairs were far more efficiently repressed by RNAi in *D. simulans* testis than in *D. melanogaster* (Figures 3B and 3C). For example, 360 of 992 siRNA-producing cis-NAT genes in *D. simulans* were significantly (FDR<0.05) derepressed in *dcr2* mutants (Table S3). Yet only 5 of 694 cis-NAT genes that generated siRNAs were derepressed in *D. melanogaster dcr2* testis (Table S3). Notably, the 5 cis-NAT mRNAs derepressed in *D. melanogaster* were distinct from the 360 cis-NAT genes derepressed in *D. simulans dcr2* testis. Genes implicated in supporting male germline development, including meiotic division, DNA repair, Golgi transport, and mitochondrial translation were overrepresented among the 360 cis-NAT mRNAs repressed by RNAi in *D. simulans* (Figure S3A).

We can envision two distinct mechanisms by which cis-NAT genes could be repressed by the RNAi pathway. First, Dicer-2 cleavage could consume mRNAs during siRNA biogenesis.^7^ Second, cis-NAT transcripts may be cleaved by siRNA-guided Ago2, because cis-NAT siRNAs are perfectly complementary to one of the two the mRNAs from which they are produced. To assess the contribution of each of these sequential steps in the RNAi pathway, we compared the degree of derepression of cis-NAT genes in *D. simulans dcr2* and *ago2* mutants. Most transcripts increased in abundance to a similar extent in *dcr2* and *ago2* testis (Spearman’s ρ = 0.75; *p* = 0; Figures 3D), suggesting that siRNA-guided Ago2-catalyzed cleavage is the main mechanism of cis-NAT gene repression by RNAi.

Dicer-2 cleavage potentially has a negligible impact on cis-NAT mRNA abundance due to the lower probability of formation of *inter*molecular dsRNAs from the complementary mRNA pairs. By contrast, hybridization of *intra*molecular duplexes in hpRNAs should be accelerated by the proximity of complementary strands. Supporting this idea, the median siRNA-to-transcript ratio for cis-NAT gene pairs was lower than that for hpRNAs (∼20-fold in *D. melanogaster*; *p* = 4 × 10^−12^; two-tailed Mann-Whitney test; ∼90-fold in *D. simulans*; *p* = 2 × 10^−25^; Figure S3B).

Conversely, siRNA-programmed Ago2 is efficient at repression of cis-NAT mRNAs likely due to its extraordinary stability and capacity for rapid target finding and multiple-turnover cleavage. Consistent with siRNA steady-state level being an important determinant of cleavage efficacy (see below), cis-NAT genes that produced abundant siRNAs were repressed to a greater degree (Figure 3B).

Because RNAi efficiency is greater in *D. simulans*, cis-NAT gene homologs in the two species are subject to distinct degrees of repression. Such divergent extents of regulation by siRNAs may result in different steady-state levels of orthologous genes in *D. simulans* vs *D. melanogaster* testis. We identified 58 genes that were efficiently repressed by RNAi in *D. simulans* (>2-fold derepression in *dcr2* testis; Table S3) but not in *D. melanogaster* (Table S3). The abundance of most mRNAs (45 of 58) was similar (<3-fold difference) in the two species (Spearman’s ρ = 0.74; Figure 3E), possibly because of the selective pressure to maintain expression level. However, the steady-state levels of 12 of the 58 mRNAs were >3-fold lower and the abundance of only one mRNA was higher in *D. simulans*, compared to *D. melanogaster* (Figure 3E). These data demonstrate the influence of siRNA-directed repression on mRNA abundance in *D. simulans*. Together, our analyses revealed the greater impact of endo-siRNAs on steady-state levels of cis-NAT gene pairs in *D. simulans*, compared to *D. melanogaster* testis.

### Rules for efficient siRNA-directed repression in vivo

*Drosophila* Ago2 is a member of a specialized AGO subclade acting in the invertebrate antiviral response.^46^ Target cleavage by fly Ago2 has been extensively studied in vitro.^47–54^ However, the in vivo parameters of efficient cleavage by endo-siRNAs have not been examined to date, in part due to the modest effects of endo-siRNAs in *D. melanogaster*. Our genetic data from testis, a dominant location of RNAi activity in *Drosophila*,^20,27^ allowed investigation of the molecular requirements for Ago2-directed repression in vivo.

Similar to piRNAs and miRNAs,^55–59^ siRNA abundance dictated the efficiency of target repression. Transcripts complementary to more abundant siRNAs were derepressed to a greater degree in *dcr2* mutant testis (Figure 4A). Importantly, the efficacy of repression correlated better with the total abundance rather than the total number of distinct siRNA sequences targeting an RNA (Figure 4A, at right).

**Figure 4.**
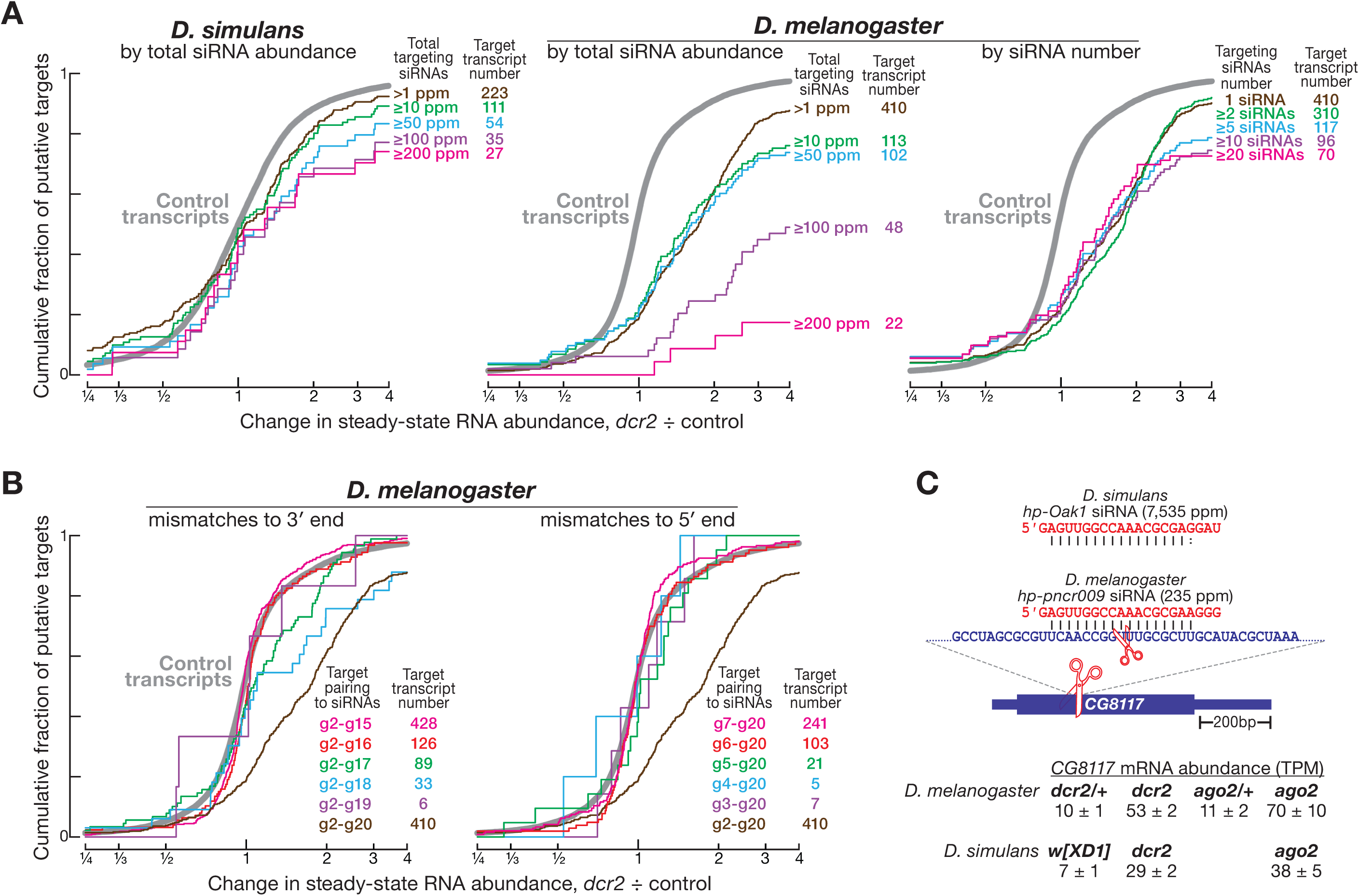
In vivo siRNA cleavage rules reveal novel targets of hpRNA-derived siRNAs in *D. simulans* testis. (A) Change in steady-state abundance of putative siRNA targets in *dcr2* vs control testis in *D. melanogaster* (*n* = 3) and *D. simulans* (*n* = 2). Data are for perfectly paired targets of siRNAs of different abundance or total siRNA number. *P* values for two-tailed Kolmogorov-Smirnov test are shown. (B) Change in steady-state abundance of putative siRNA targets in *dcr2* vs control testis in *D. melanogaster* (*n* = 3). Data are for perfectly and partially paired targets of siRNAs. *P* values for two-tailed Kolmogorov-Smirnov test are shown. (C) Both in *D. melanogaster* and *D. simulans*, *CG8117* mRNA is targeted by an abundant hpRNA-derived siRNA via g2-g18 complementarity. Mean ± SD of steady-state abundance in *D. melanogaster* (*n* = 3) and *D. simulans* (*n* = 2) controls and RNAi mutants is shown.

To examine whether mismatches between target and guide are tolerated by Ago2, we identified siRNA:target pairs with unpaired nucleotides either at the siRNA 5′ or 3′ end. For stringency, we confined comparisons to the non-overlapping sets of siRNA:target pairs and excluded transcripts also targeted by perfectly complementary siRNAs. Applying these constraints was only possible for *D. melanogaster* data, owing to the sparsity of siRNAs that allowed avoiding the overlap among siRNAs with partial and full complementarity to mRNAs.

Ago2-catalyzed cleavage tolerated mismatches to siRNA 3′ nucleotides, but not its 5′ sequence (Figure 4B). Derepression of mRNAs in *dcr2* was detected for siRNA:target pairs with mismatches between nucleotides g18–g20. In fact, siRNAs paired to targets with just nucleotides g2–g17 (16 nt) conferred detectable repression (Figure 4B, at left). Conversely, mRNA steady-state levels were unaltered in *dcr2* mutant testis for transcripts with mismatches to 1–5 nt of siRNA 5′ sequence. Even a single seed mismatch at siRNA nucleotide g2 prevented repression by Ago2 (Figure 4B, at right). These results agree with in vitro cleavage experiments showing that Ago2 tolerates mismatches near the 3′ end of the guide.^50,54^ We conclude that efficient repression by Ago2 is promoted by high siRNA abundance and requires perfect pairing to the siRNA seed and central sequence.

### Novel targets of hpRNA-derived siRNAs

Using the minimal requirement of g2–g17 pairing (Figure 4B), we identified 535 putative siRNA targets in *D. melanogaster*, including 461 transposons and 74 mRNAs and lncRNAs. Of the 535 putative targets, 61 were significantly (FDR<0.05) derepressed in *dcr2* testis (Table S4). Consistent with the important role of *D. melanogaster* siRNAs in silencing transposons, 60 of the 61 derepressed targets were derived from DNA transposons and retroelements. One mRNA target—*CG8117*, the fly ortholog of human Transcription elongation factor A N-terminal and central domain containing (TCEANC)—was also significantly derepressed in *dcr2* mutants (Figure 4C). While transposon loci were targeted by one or many siRNAs of low abundance (∼1–2 ppm), *CG8117* mRNA was repressed by a single abundant (∼235 ppm) siRNA (Table S4) derived from *hp-pncr009*. This siRNA base paired with *CG8117* from g2 to g18 and repressed the target mRNA by ∼5-fold (Figure 4C).

Argonaute proteins cleave complementary transcripts at the phosphodiester bond joining the target nucleotide (t10 and t11) paired to siRNA nucleotides g10 and g11. Argonaute cleavage leaves a 5′-monophosphate, producing a monophosphorylated RNA whose 5′ end pairs with siRNA nucleotides g2–g10 (Figure S1D). We used 5′-monophosphate RNA-specific sequencing to detect cleavage products generated by siRNA-directed slicing in *D. melanogaster* testis. Consistent with the *hp-pncr009* siRNA guiding cleavage of *CG8117* mRNA, the 5′-monophosphorylated RNA fragment, predicted to be generated by Ago2, was present in the control but reduced >30-fold in *dcr2* testis (control: 0.07 ± 0.02 ppm, *n* = 3; *dcr2*: 0.002 ± 0.002 ppm, *n* = 3; *p* = 10^−22^, unpaired, two-tailed Mann-Whitney test). However, cleavage fragments from transposon transcripts were undetectable, potentially due to the low abundance of repeat-targeting siRNAs and transposon transcripts and therefore low steady-state levels of transposon-derived cleavage products. Resequencing 5′-monophosphate-bearing RNA libraries to a depth of one billion reads each did not improve the sensitivity of cleavage fragment detection. The rapid XRN1-catalyzed decay of 5′ monophosphate-bearing RNAs in the cytoplasm may prevent detection of low-abundance cleavage products.

In *D. simulans*, of 417 putative targets, 95—including 7 previously reported—were significantly (FDR<0.5) derepressed in *dcr2* testis (Table S5). Notably, 32 of the 95 mRNAs were targeted by abundant siRNAs (>10 ppm; Table S5). Of these 32 high-confidence mRNA targets, 19 were fully complementary to siRNAs and 13 paired with siRNAs via nucleotides g2–g17, g2–g18, or g2–g19 (Table S5).

TCEANC (CG8117) is the only gene targeted by an siRNA *in trans* in *D. melanogaster*. *D. simulans* produces an extremely abundant (∼7,535 ppm) hpRNA-derived siRNA complementary to the same 17-nt site in DsTCEANC. In fact, the location of the cleavage site in the two species was identical. Remarkably, the sequences of the two siRNAs differ only at position g18, which makes an A:U base pair in *D. melanogaster* and a G:U wobble pair in *D. simulans*, despite an overall lack of conservation of their respective hpRNA precursors (Figure 4C). We conclude that, in addition to the control of selfish meiotic drives, hpRNA-derived siRNAs repress dozens of host mRNAs in *D. simulans* testis.

## DISCUSSION

### Molecular determinants of siRNA biogenesis

Our analyses demonstrated that Ago2 is required for maturation of perfectly paired siRNA duplexes. Choice of siRNA guide was only modestly influenced by the identity of 5′ terminal nucleotide in *D. simulans* and virtually every siRNA duplex yielded at least one siRNA strand (Figure 2C).

### Rules for efficient siRNA targeting in vivo

We and others have recently demonstrated that PIWI-clade Argonaute proteins tolerate target mismatches to piRNA seed and nucleotides adjacent to the scissile phosphate.^58,60^ However, the requirements for functional repression by endo-siRNAs have so far remained undefined. Previous biochemical experiments suggest that siRNAs can cleave targets via imperfect complementarity, yet it was unclear whether partial pairing can mediate productive repression *in vivo*.^54^ Our analyses demonstrate that target mismatches to the siRNA seed and central nucleotides are not tolerated *in vivo* (Figure 4B). We also find that the high abundance of siRNAs predicts productive target repression (Figure 4A). Incidentally, the steady-state levels of target cleavage products produced by fly siRNAs were much lower than those generated by fly and mouse piRNAs.^58,61–63^ Perhaps (1) slow product release by PIWI proteins stabilizes cleavage fragments generated by piRNAs^64,65^ or (2) the decay pathways for cleaved targets operate differently between cell types or mammals and insects.

### mRNA repression by endo-siRNAs is required for *D. simulans* spermatogenesis

Our data show that the breadth of siRNA-directed gene repression extends far beyond the handful of hpRNA loci identified by studies of *Drosophila* RNAi biology to date. siRNA steady-state levels are tenfold higher in *D. simulans* than in *D. melanogaster* testis, permitting productive repression by *D. simulans* siRNAs of multiple endogenous mRNAs via two mechanisms. First, hundreds of overlapping cis-NAT genes generate abundant siRNAs that cleave mRNAs from which they were produced (regulation *in cis*; Figure 3). Second, hpRNA-derived siRNAs repress dozens of *D. simulans* mRNAs *in trans*, via partial complementarity (Figure 4). *D. simulans* RNAi mutants are fully sterile, due to massive defects in spermatogenesis.^20^ Because many targets of *D. simulans* siRNAs are implicated in supporting male germline development, we propose that siRNA-directed repression of host mRNAs is required for successful spermatogenesis in *D. simulans*.

### Evolutionary flux of endo-siRNA targets

Repression of endogenous mRNAs is likely an evolutionarily derived function of RNAi—an addition to its ancestral role in antiviral defense. Similarly, the piRNA pathway silences transposons in nearly all animals, yet an evolutionarily derived piRNA class, the pachytene piRNAs, also regulates host genes during spermatogenesis.^63,66–72^

The distinct repertoires of siRNA-target networks in *D. melanogaster* and *D. simulans*, species whose genomes are nearly identical, highlight the extreme evolutionary lability of the endogenous RNAi pathway. Gene repression by endo-siRNAs has largely been unknown due to their marginal impacts in *D. melanogaster*. Beyond our proposal that hpRNAs are frequently utilized to suppress selfish meiotic drivers^24,25^, we find that numerous other testis genes are subject to species-specific control by RNAi. These results suggest the existence of additional layers of rapidly evolving intragenomic conflict and underscore the need to characterize and manipulate the RNAi pathway across more model systems in the future.

## ACKNOWLEDGMENTS

We thank the Zegar Family Foundation for their generous support; staff at the NYU Center for Genomics and System Biology Genomics Core for their assistance and resources; Phillip Zamore, members of the Gainetdinov and Lai laboratories for critical comments on the manuscript. Work in the ECL lab was supported by the National Institute of Child Health and Human Development (R01-HD108914), National Institute for General Medical Sciences (R01-GM083300) and the National Cancer Institute (MSK Core Grant P30-CA008748). Work in the JS lab was supported by the National Science Foundation (NSF 2427326).

## SUPPLEMENTARY FIGURE TITLES AND LEGENDS

**Figure S1.**
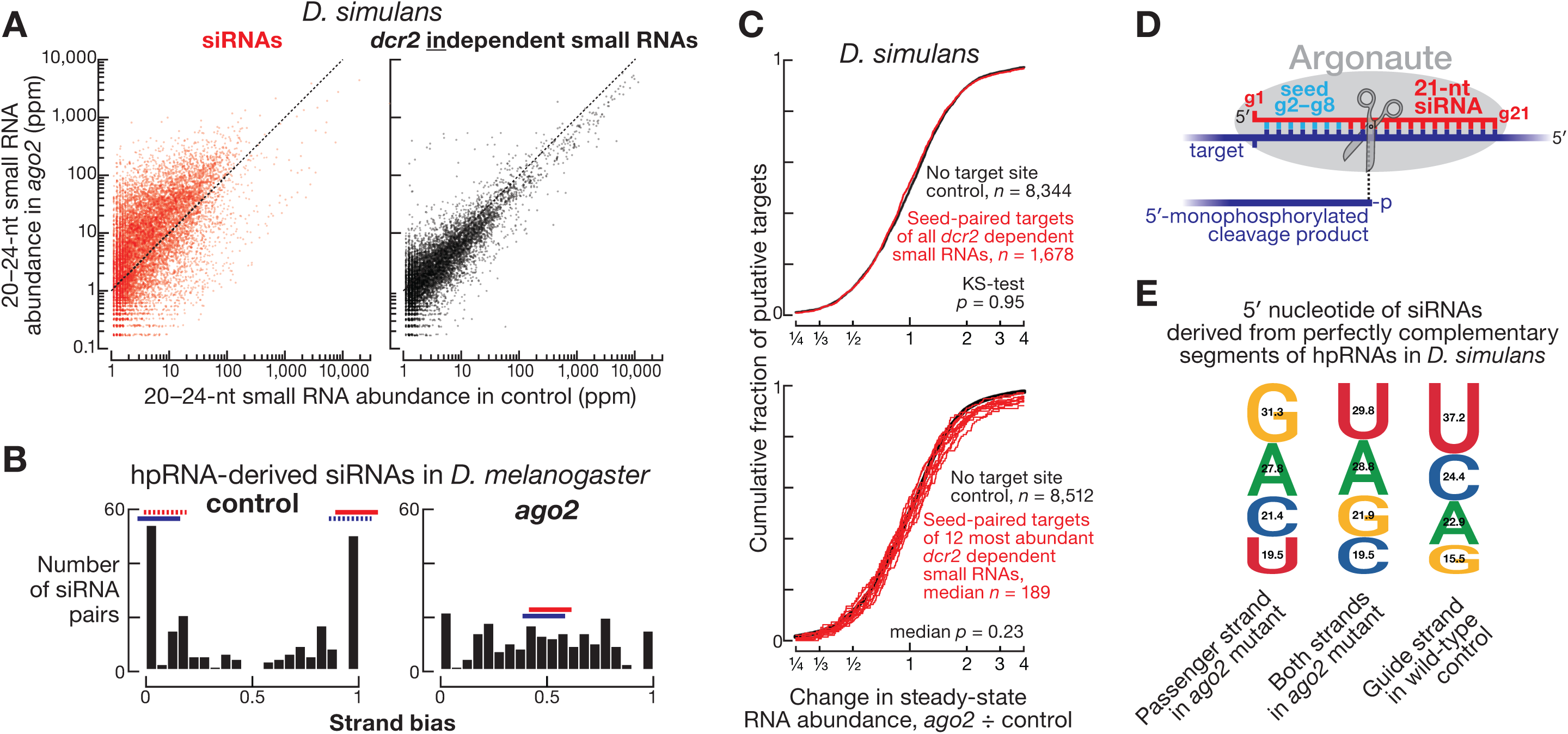
In *ago2* mutants, siRNAs persist as double-stranded intermediates, Related to Figure 2. (A) Mean (*n* = 2) abundance of ≥1 ppm, 20–24-nt small RNAs in control and *ago2* testis in *D. simulans*. Abundance of 20–24-nt small RNAs with the same 5′ prefix was summed. *dcr2*-dependent siRNAs are defined as decreased by ≥10-fold or undetectable in *dcr2* testis and are shown in red; all other small RNAs are assigned as *dcr2*-independent and are shown in black (see Figure 1A). (B) Mean (*n* = 2) strand bias was calculated as the abundance of top strand divided by the sum of abundances of top and bottom strands. Data are from control and *ago2* testis for perfectly complementary duplexes derived from hpRNA loci in *D. melanogaster*. (C) Change in abundance in *ago2* mutant vs control for mRNAs paired via nucleotides g2–g8 to all (upper panel) or 12 most abundant siRNAs in *D. simulans* (lower panel). siRNAs are the small RNAs in Figure 1A defined as *dcr2*-dependent. *P* values for two-tailed Kolmogorov-Smirnov test are shown. (D) siRNAs direct Argonaute proteins (e.g., Ago2) to cleave complementary targets. (E) Identity of 5′ terminal nucleotide for both strands of siRNA duplexes or only passenger strand in *ago2* testis and for guide strand in wild-type control. Guide strands were defined as siRNAs whose abundance was unchanged or decreased <2-fold in control vs *ago2*. Passenger strands were defined as siRNAs whose abundance decreased >10-fold in control vs *ago2*. Data are for siRNAs derived from perfectly complementary segments of hpRNAs in *D. simulans*.

**Figure S2.**
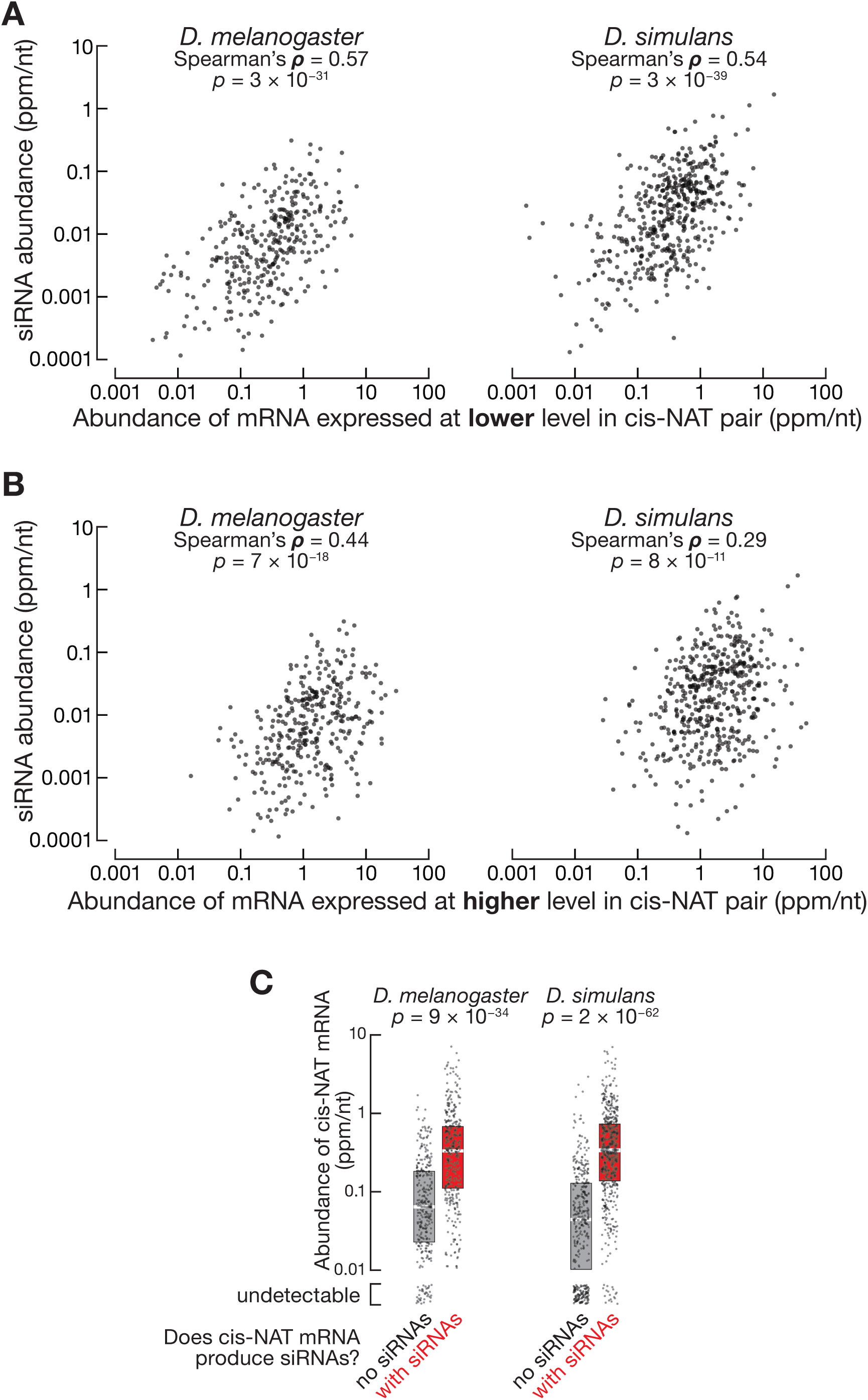
dsRNA substrate abundance dictates cis-NAT siRNA levels, Related to Figure 3. (A) Mean coverage of cis-NAT siRNAs and cis-NAT transcripts normalized to the length of cis-NAT overlap in *D. melanogaster* (*n* = 3) and *D. simulans* (*n* = 2). Data are for all siRNAs from each cis-NAT and transcripts with the lower abundance in the cis-NAT pair. Rank-order Spearman’s correlation coefficient and *p* value are shown. (B) Mean coverage of cis-NAT siRNAs and cis-NAT transcripts normalized to the length of cis-NAT overlap in *D. melanogaster* (*n* = 3) and *D. simulans* (*n* = 2). Data are for all siRNAs from each cis-NAT and transcripts with the higher abundance in the cis-NAT pair. Rank-order Spearman’s correlation coefficient and *p* value are shown. (C) Mean coverage of cis-NAT transcripts normalized to the length of cis-NAT overlap in *D. melanogaster* (*n* = 3) and *D. simulans* (*n* = 2). Data are for cis-NAT transcripts that produce or do not siRNAs (Table S3). *P* value for unpaired, two-tailed Mann-Whitney test are shown.

**Figure S3.**
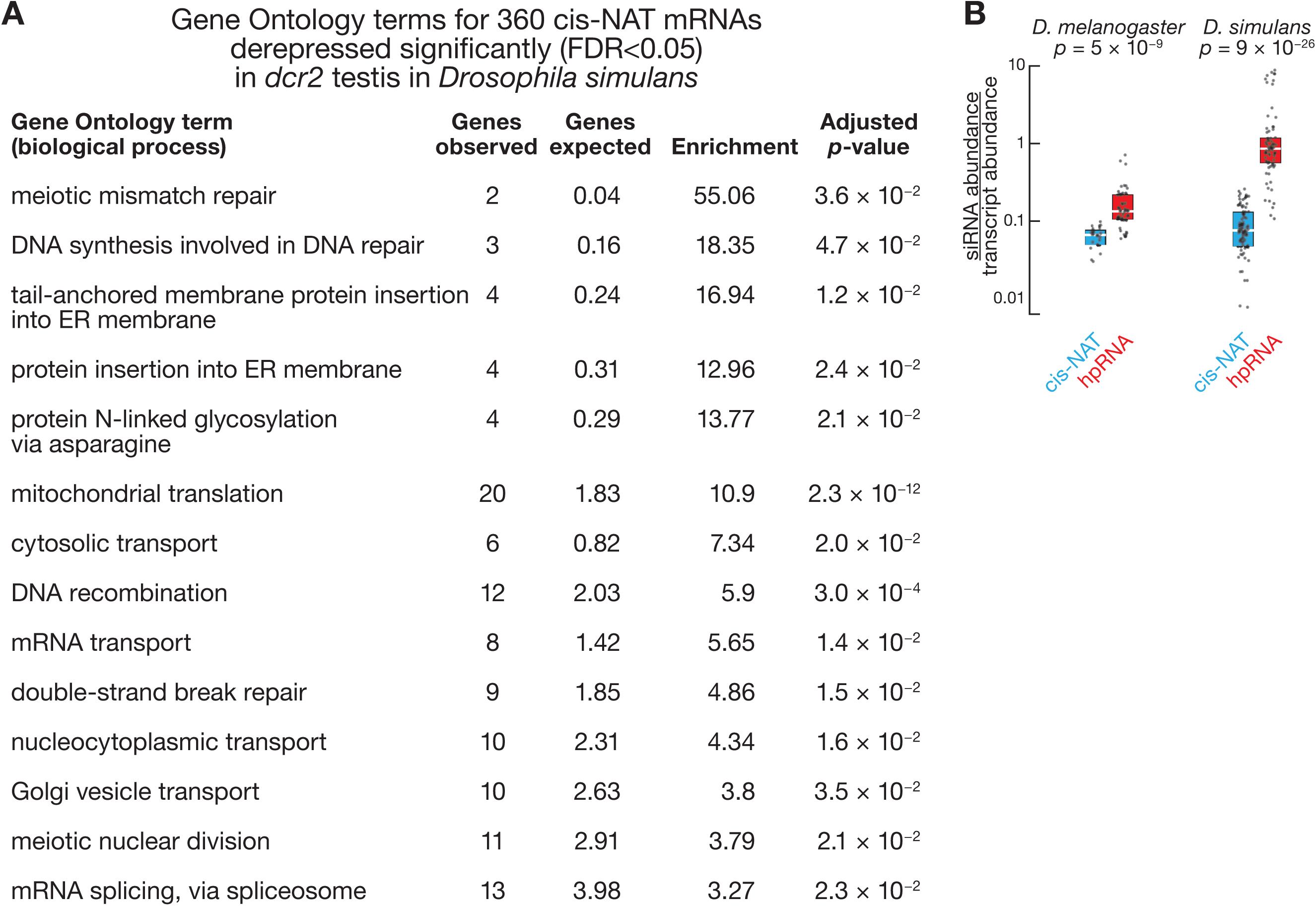
cis-NAT siRNAs repress hundreds of mRNAs in *D. simulans* testis, Related to Figure 3. (A) Gene ontology terms enriched among 360 cis-NAT mRNAs repressed by RNAi in *D. simulans*. Enrichment for Gene Ontology terms (biological processes) was calculated using Panther database with two-tailed Fisher’s exact test and *p* value adjusted for multiple comparisons with Benjamini-Hochberg procedure. (B) Mean coverage of siRNAs normalized to mean coverage of source transcripts in *D. melanogaster* (*n* = 3) and *D. simulans* (*n* = 2). Data are for siRNAs from cis-NAT and hpRNA loci. *P* value for unpaired, two-tailed Mann-Whitney test are shown.

